# Using GIS Dashboards to highlight AMR data disparities in Africa for Policy, Research, and Public Health

**DOI:** 10.64898/2026.08.09.743821

**Authors:** Wendy Akushika Dogbegah, Stephen Obol Opiyo, Peace Proscovia Aber, Christian Kaembou Tiambo

## Abstract

Antimicrobial resistance (AMR) continues to pose a major public health threat across Africa, yet available surveillance data remain highly fragmented across private, public, and academic sources. This study analysed continent-wide AMR surveillance patterns by integrating datasets from multiple independent repositories and visualising them through interactive Geographic Information System (GIS) dashboards. The objective was to generate an integrated evidence base that highlights resistance patterns, surveillance disparities, reporting gaps, and opportunities for improved AMR monitoring across Africa. Data were compiled from major private AMR surveillance programmes including Pfizer’s ATLAS, GSK’s SOAR, Johnson & Johnson’s DREAM, Venatorx’s GEARS, and Shionogi’s SIDERO-WT covering the period 2004–2022. Public datasets from the WHO Global Antimicrobial Resistance and Use Surveillance System (GLASS) and the Fleming Fund’s Mapping Antimicrobial Resistance and Antimicrobial Use Partnership (MAAP) were incorporated for 2016–2020, together with published AMR studies conducted between 2010 and 2024. Datasets were harmonised to align key variables including bacterial species, isolate identifiers, antibiotics tested, surveillance source, geographical location, and categorical AMR outcomes while preserving the original structure of the contributing datasets. Interactive dashboards were developed using R Shiny to support spatial visualisation and dynamic analytical exploration of resistance patterns, temporal trends, species distribution, and country-level surveillance coverage. Descriptive analyses including means, standard deviations, medians, interquartile ranges (IQR), frequency distributions, Gini coefficients, Shannon entropy, Herfindahl–Hirschman Index (HHI), and Lorenz curves were used to assess inequality and concentration in country-level AMR reporting across surveillance systems. The integrated analyses revealed substantial heterogeneity and concentration in AMR surveillance reporting across Africa, reflecting major differences in surveillance intensity, laboratory infrastructure, reporting systems, and diagnostic capacity across countries. Private datasets demonstrated broader antibiotic panels and longer temporal coverage, whereas public datasets exhibited substantial gaps in country participation and pathogen-antibiotic representation. Published AMR studies additionally highlighted important surveillance information absent from formal surveillance databases. By integrating multiple streams of AMR evidence, this study demonstrates the value of interactive GIS dashboards as exploratory and updateable surveillance-support tools for improving visibility of fragmented AMR datasets, identifying surveillance disparities, supporting geographically informed interpretation of resistance trends, and strengthening future AMR surveillance harmonisation efforts across Africa.

## Introduction

Antimicrobial resistance (AMR) has rapidly emerged as one of the most critical challenges facing global public health in the 21st century.^1^ The growing prevalence of drug resistant pathogens threatens to undermine decades of medical progress, as treatments that were once highly effective now struggle to combat infections.^2^ This rise in resistance is not limited to a specific region or demographic but spans across continents, affecting both developed and developing countries alike. The consequences are stark: infections that were once easily treatable now result in prolonged illnesses, higher mortality rates, and an increasing burden on healthcare systems.^3^ The escalating presence of AMR exacerbates medical complexities and necessitates longer hospital stays, placing significant pressure on already strained healthcare resources worldwide.^4^

Addressing AMR requires urgent, coordinated efforts across various sectors. The medical community must improve diagnostics and develop more targeted treatment protocols, while policymakers must prioritise public health strategies that emphasise enhanced surveillance and data collection.^5^ However, these efforts are often hindered by the fragmented nature of AMR data, especially in low and middle income countries (LMICs) where data collection infrastructures are insufficient.^5^ In Africa, the scarcity of reliable and comprehensive AMR data is a significant barrier to effective intervention, with many countries struggling to maintain consistent and up-to-date surveillance.^6,7,8^

Fragmented AMR surveillance systems have major consequences for clinical care, public health decision-making, and health system preparedness. In many LMICs, incomplete laboratory reporting, inconsistent data sharing, and limited integration between surveillance platforms delay the detection of emerging resistance trends and reduce the ability of health authorities to respond rapidly to outbreaks of resistant pathogens^6,7,8^. Delayed recognition of resistance patterns contributes to inappropriate antimicrobial prescribing, increased treatment failure, prolonged hospitalization, and higher mortality rates among infected patients^1,3^. In addition, fragmented surveillance systems limit the ability to identify geographical hotspots of resistance and weaken the implementation of evidence-based antimicrobial stewardship programmes and infection prevention strategies^4,5^.

The economic and operational burden associated with fragmented AMR surveillance is also substantial. Weak integration of surveillance systems can lead to inefficient allocation of limited laboratory infrastructure, diagnostic resources, and public health funding, particularly in resource-constrained African settings^5,6^. Countries with incomplete or inconsistent AMR reporting may struggle to prioritise high-risk pathogens, monitor the effectiveness of interventions, or allocate resources to regions with the greatest surveillance needs^7,8^. Furthermore, fragmented data systems hinder continental and global coordination efforts aimed at combating AMR and may reduce the effectiveness of regional One Health surveillance strategies^4^. Strengthening integrated and geographically informed AMR surveillance systems is therefore essential for improving public health preparedness, supporting timely interventions, and reducing the long-term clinical and economic burden associated with antimicrobial resistance.

A key issue in Africa’s AMR data crisis is the disparity between public and private data sources. Publicly available datasets, such as those from the World Health Organisation’s Global Antimicrobial Resistance Surveillance System (GLASS)^9^, and the African Network for Drugs and Diagnostics Innovation ^10^, The Mapping Antimicrobial Resistance and Antimicrobial Use Partnership (MAAP;^11^), are often aggregated at a high level, lacking the resolution necessary for localised interventions. These datasets typically focus on broader trends, which can obscure critical insights needed for targeted public health responses. In contrast, private sector data, particularly from pharmaceutical companies, often contains detailed information on drug resistance patterns, but much of this data remains inaccessible to public health authorities and researchers.^12^ This disconnect between public and private data sources hampers a comprehensive understanding of AMR trends, particularly in regions where healthcare systems are fragile.

The integration of public and private AMR data is, therefore, essential to developing a more complete and accurate picture of resistance patterns in Africa. By combining these datasets, this study seeks to bridge the current data gaps, enabling more effective surveillance and intervention strategies. Moreover, the study proposes the innovative use of Geographic Information System (GIS) mapping to overlay AMR data onto geographical regions across Africa. GIS technology has been extensively used in infectious disease control, providing critical insights into disease spread and resource allocation.^13^ However, its application in tracking AMR has been limited, particularly in Africa. By employing GIS mapping in this context, the study aims to visualise the geographic spread of resistance, identify potential hotspots, and support more localised and effective public health interventions.

The dashboards developed in this study are intended to complement, rather than replace, existing antimicrobial resistance surveillance frameworks operating at global, regional, and national levels. Specifically, the integrated visualisation approach aligns with the objectives of the World Health Organization (WHO) Global Action Plan on Antimicrobial Resistance, which emphasises strengthening surveillance and research through coordinated data collection and sharing. The dashboards also support the objectives of the Global Antimicrobial Resistance and Use Surveillance System (GLASS) and national AMR National Action Plans (NAPs) implemented across African countries by providing an additional framework for integrating, visualising, and interpreting AMR data from multiple surveillance sources. Furthermore, the dashboards are consistent with One Health surveillance approaches that promote coordinated monitoring of antimicrobial resistance across human, animal, food, and environmental sectors^4,7^. By improving visibility of reporting gaps, temporal trends, and geographical disparities across surveillance systems, the dashboards may support surveillance coordination, data interpretation, and regional AMR monitoring activities while preserving the original structure and ownership of contributing datasets.

In addition to integrating public and private datasets, this study will also pilot incorporating country specific academic published research on AMR into a comprehensive, interactive dashboard. Many of these studies, though crucial, remain absent from formal AMR surveillance databases, both public and private. The primary objective is to introduce the AMR research community to a source of AMR data from Africa that has not yet been incorporated into a database and to encourage researchers who publish their data to contribute this information to the pilot project. By highlighting and visualising this academic research data, the dashboard will offer a valuable resource for researchers, policymakers, and healthcare providers, enhancing the accuracy of AMR surveillance and facilitating evidence based decision making.

In summary, this study aims to integrate diverse public and private datasets, incorporating country specific academic research data, and utilising GIS dashboards. This holistic approach is designed to enhance Africa’s ability to monitor and respond to AMR effectively, improve the impact of public health interventions, and ensure optimal allocation of resources. Ultimately, our study seeks to advance the development of more effective strategies to combat AMR, thereby safeguarding the health and wellbeing of populations both within Africa and beyond.

## Methods

### Ethical Considerations

This study utilises data sourced from both private and public sectors, as well as AMR data published in academic papers. Approval for the use of private data was obtained under the approval number AMR ID: 00010209 from Vivli.^14^ The public data was acquired from GLASS^9^, and MAAP.^11^ The data from published papers were gathered from two review papers. ^15,16^ Ethical approval and individual informed consent were not required for this study.

Data governance and privacy considerations were incorporated throughout the data acquisition and integration process. Certain datasets used in this study, particularly private-sector surveillance resources such as ATLAS and related Vivli-hosted datasets, were obtained through controlled-access approval mechanisms under the Vivli antimicrobial resistance data-sharing framework^14^. Only aggregated and derived analytical outputs were publicly deposited and visualised within the dashboards. No individual-level patient identifiers, personal health information, or directly identifiable clinical records were included in the integrated analytical datasets or dashboard visualisations. Use of all datasets complied with the approval conditions, governance requirements, and data-sharing restrictions established by the contributing surveillance systems, pharmaceutical partners, and public health platforms.

### Data Sources

#### Private data

Pharmaceutical companies gather extensive drug resistance data during their clinical trials and drug development processes. Data from Pfizer (ATLAS; 2004 to 2022), GSK (SOAR; 2015 to 2018), Johnson & Johnson (DREAM; 2011 to 2019), Venatorx (GEARS; 2018 to 2020), and Shionogi (SIDERO-WT; 2013 to 2016) were obtained from the Vivli database.^12,14^ These datasets were downloaded on June 26th, 2024. We extracted information on African countries from each dataset, including the year AMR tests were conducted, the country where the tests took place, the most commonly used antibiotics in Africa, isolate IDs, species, and phenotypes. However, not all datasets contain every piece of this information.

#### Public data

The GLASS dataset, acquired from Rahbe and colleagues^17,9^, covers the period 2017–2020 and includes the following pathogens: *Acinetobacter* spp., *E. coli*, *Klebsiella pneumoniae*, *Neisseria gonorrhoeae*, *Salmonella* spp., *Shigella* spp., *Staphylococcus aureus*, and *Streptococcus pneumoniae*.^9^ We extracted information on African countries, including the year AMR tests were conducted and the AMR susceptibility tests on June 26th, 2024. MAAP reviewed 819,584 AMR records from 2016 to 2019, collected from 205 laboratories across 14 African countries.^11^ We extracted the counts of resistance from the reports of MAAP, specifically “Table 8: AMR Estimates for WHO priority pathogens” across all 14 countries on June 26th, 2024, available at the African Society for Laboratory Medicine.^18^

#### ATLAS data

To provide a comprehensive in-depth of AMR data across all African nations, we plan to develop a dashboard from a private dataset. The ATLAS dataset^12^, which includes data from 13 African countries collected between 2004 and 2022, will be a key resource for this initiative. From this dataset, we have extracted vital information on bacterial species, countries, sample sources, sample counts, gender, and antibiotics. This dashboard will consolidate and present the data in a clear, accessible manner, enhancing the ability to analyse and interpret AMR trends across the continent.

The private ATLAS dataset spans a substantially longer surveillance period (2004–2022) compared to the public datasets, including GLASS and MAAP, which primarily cover the period from approximately 2016 to 2020. Consequently, the differences observed in isolate counts, country participation, and temporal coverage should not be interpreted as direct comparative measures of antimicrobial resistance burden or surveillance performance between private and public surveillance systems. Rather, the dashboards were designed primarily to describe and visualise AMR surveillance coverage, reporting trends, isolate distribution, and spatial representation across available African datasets while preserving the original structure of the contributing data sources.

The observed disparities in isolate counts and reporting coverage likely reflect differences in surveillance duration, reporting intensity, laboratory infrastructure, diagnostic capacity, participation of reporting laboratories, and overall data availability across surveillance systems. To improve transparency, annual country-level reporting summaries and per-year isolate count descriptions were included in the revised manuscript and supplementary materials to demonstrate fluctuations in surveillance participation over time. Raw isolate counts are influenced by surveillance intensity and should therefore not be interpreted as direct estimates of disease burden or AMR prevalence across countries. Although normalization approaches such as isolates per million population or adjustment by reporting laboratories could improve comparability, consistent metadata on laboratory participation and standardized denominators were not uniformly available across all datasets and countries, limiting the feasibility of applying harmonized normalization methods across the full surveillance period. These limitations are important considerations for future AMR surveillance harmonization studies.

#### Research published papers data

Our inclusion criteria for AMR research papers required that the selected studies cover a majority of countries that were either missing or underrepresented in both private^12^ and public^9,11^ databases and report the percentage of resistance to the most commonly used antibiotics in African countries. We chose two review articles, da Costa RC et al.^15^ and Gahimbare L et al.^16^, as they employed rigorous inclusion and exclusion criteria when selecting published AMR studies. The countries selected from these two papers for inclusion in the pilot dashboard project were Algeria^19^, Angola^20,21,22,23^, Benin^24^, Central African Republic^25^, Democratic Republic of Congo^26^, Egypt^27^, Eritrea^28^, Ethiopia^29^, Ghana^30^, Mauritius^31^, Mozambique^32^, South Africa^33^, Tanzania^34^, and Uganda^35^. The literature search was conducted on June 26^th^, 2024.

#### Non-Aggregated ATLAS Dataset Processing

Among the included AMR data sources, ATLAS was the only dataset available in both aggregated and non-aggregated formats. The non-aggregated ATLAS dataset contained isolate-level records and was processed separately from the aggregated datasets obtained from GLASS, MAAP, and other surveillance sources. Variables within the isolate-level ATLAS dataset included country, year, bacterial species, antibiotic, interpretation category, and associated metadata. Antibiotic names and susceptibility interpretation categories were retained as originally reported in the ATLAS dataset, while duplicate isolate records were assessed using available isolate identifiers and combinations of species, country, year, antibiotic, and source metadata to minimize double counting. Antibiotics that were not tested for specific isolates were labelled as “No Info” to distinguish untested antibiotics from resistant, intermediate, or susceptible results. Quality assurance procedures included verification of country names and coordinates, validation of year ranges, inspection of susceptibility interpretation distributions, and comparison of processed isolate counts against the original ATLAS dataset summaries prior to integration into the interactive dashboard and downstream analyses.

#### Identification of coordinates

To support the development of the interactive GIS dashboard, geographic coordinates were assigned to each country included in the analysis. Country-level latitude and longitude coordinates were obtained and linked to the surveillance datasets to enable spatial visualization and mapping of AMR reporting patterns across Africa.

#### Integration of private and public datasets

To support harmonization and reproducibility, supplementary materials were developed including Supplementary Table S1 (Unified Data Dictionary), Supplementary Table S2 (Variable Mapping Table), and Supplementary Table S3 (Antibiotic Harmonization Table). These materials document how variables from public and private datasets were aligned into a unified analytical schema used for integration, visualization, and downstream analysis. We extracted coordinates, isolates, countries, source of data (e.g., ATLAS, GLASS, and MAAP), classification (private or public), and the year in which the data were collected. For Figure 1, isolates were aggregated by country and year to compare the relative contribution of public and private AMR datasets across African countries (Supplementary Table S4).

**Fig 1:** Interactive GIS dashboard displaying 834,365 isolates categorized by data source (private datasets^12^: ATLAS, DREAM, GEARS, SIDERO-WT, and SOUR; public datasets^9,11^: MAAP and GLASS) and their distribution across African countries and years. Private datasets covered the period from 2004 to 2022, whereas public datasets covered the period from 2016 to 2020. The dashboard also includes a geographical map showing country-level data collection locations across Africa using country centroid coordinates. Fig 1 was generated from the Africa_AMR_Country_Year_Spatial_Data.xlsx **(Supplementary Table S1)^Error! Reference source not found.^**, which contains the country-year isolate summaries, spatial coordinates, classification variables, and surveillance source information used to construct the GIS-based spatial visualization.

We did not standardise or normalise the private and public datasets before integration, as the objective was to visualise the original reported data while preserving the structure of the source datasets. Instead, datasets were harmonized by aligning variables into a common schema, including country, year, isolate count, source, classification, and geographic coordinates. Country names were aligned across datasets, and each country was assigned unique coordinates to support geographical visualisation within the dashboard. For antibiotic-level analyses, missing values represented antibiotics that were not tested for specific isolates in the original datasets. These missing values were labelled as “No Info” in R during preprocessing to distinguish unavailable testing information from susceptibility interpretations.

Because antimicrobial susceptibility testing methodologies, breakpoint interpretation criteria, laboratory quality assurance procedures, and isolate selection practices differed substantially across surveillance systems and countries, resistance frequencies presented within the integrated dashboards should not be interpreted as directly comparable epidemiological estimates across all datasets. The contributing surveillance systems and published studies were developed independently and therefore reflected variations in clinical sampling strategies, antimicrobial susceptibility testing standards (including CLSI, EUCAST, and locally adopted laboratory protocols), reporting intensity, healthcare infrastructure, diagnostic capacity, and surveillance duration. Consequently, observed differences in resistance frequencies across countries and datasets may reflect both underlying biological resistance patterns and methodological differences in surveillance implementation and laboratory reporting practices^6,7,8^.

Rather than generating standardised continental prevalence estimates, the primary objective of integrating these datasets was to support descriptive visualisation, exploratory analysis, and assessment of AMR surveillance representation across African countries and surveillance systems. The dashboards were therefore designed to highlight surveillance coverage, reporting disparities, temporal trends, geographical distribution of isolates, and broad pathogen-antibiotic resistance patterns while preserving the original structure of the contributing datasets. Interpretation of resistance summaries within the dashboards should therefore be performed cautiously and within the context of the substantial heterogeneity in laboratory methodologies, reporting practices, isolate inclusion criteria, and surveillance capacities across the contributing datasets. This study consequently emphasises surveillance visibility and exploratory comparative assessment rather than direct epidemiological comparison of resistance prevalence between countries or surveillance systems.

#### Development of GIS web-based dashboard from integrated public and private datasets

The dashboard of integrated data was developed using R Shiny^36-**Error! Reference source not found.**^. This cloud hosted dashboard is built on DigitalOcean^38^ platform and rely on datasets containing latitude and longitude coordinates.

#### Development of GIS web-based dashboard from ATLAS data

We developed a dashboard from ATLAS dataset from Pfizer to detail information on bacteria, antibiotics, source of samples since it has data from Africa since 2004 using R Shiny^36–37^.

#### Development of pilot research published dashboard using R Shiny Dashboard

This dashboard was created using the R Shiny dashboard^36, 38^ platform, which allows for an interactive and user friendly interface. Users interact with the dashboard through customisable filters, selecting specific country, and species, to tailor the analysis to their needs. The Shiny platform’s capabilities enable dynamic visualisation, ensuring that the data is both accessible and interpretable.

#### Descriptive statistics

The aim of this study is to present AMR datasets from both private and public databases through a dashboard visualisation. To maintain the integrity of the original data, we did not impute missing values. Instead, we focused on computing basic descriptive statistics, including total counts, mean, standard deviation, median, interquartile range, and percentage within the respective databases. The calculations were done in R version 4.4.0.^37^

#### Inequality and concentration analysis

To assess the degree of imbalance and concentration in country-level AMR isolate contributions across surveillance datasets, inequality and concentration metrics were calculated for both the public and private datasets. Metrics calculated included the Gini coefficient, Shannon entropy, normalized entropy, Herfindahl–Hirschman Index (HHI), and Lorenz curves. The Gini coefficient was used to quantify inequality in isolate distribution across countries, where values closer to 1 indicate stronger concentration and inequality. Shannon entropy and normalized entropy were used to assess the diversity and distributional evenness of isolate contributions across countries, while the HHI was used to measure the degree of concentration in surveillance reporting among dominant contributing countries. Lorenz curves were additionally generated to visually illustrate deviations from equal isolate distribution across countries. The analyses were conducted in R version 4.4.0^37^.

## Results

### Distribution of AMR data across Africa from private and public databases

Table 1 presents the number and proportion of isolates from public datasets (MAAP and GLASS) across 22 countries between 2016 and 2020, highlighting significant disparities in data collection efforts. South Africa recorded the highest number of isolates at 488,511, accounting for 60.87% of the total, followed by Tunisia with 158,960 isolates (19.81%) and Uganda with 25,316 isolates (3.15%). In contrast, countries like Sierra Leone, Mozambique, and Sudan reported substantially lower isolate counts, representing only 0.01%, 0.04%, and 0.12% of the total, respectively. The overall mean number of isolates per country was 36,480, with a substantial standard deviation of 106,148, reflecting the wide variation in isolate counts across different countries. These findings underscore the uneven distribution of AMR data collection efforts across Africa, with some countries contributing significantly more to surveillance than others. These results are similar to the studies reported by. ^6,7,8^

**Table 1:** The table summarizes AMR data collected from public databases (GLASS^9^ and MAAP^11^) across 22 African countries. It includes the number of isolates for each country and provides the mean and standard deviation of AMR data from 2016 to 2020, as detailed in the table.

| No | Country | Total isolates | Percentage | Year |
| --- | --- | --- | --- | --- |
| 1 | Burkina Faso | 7,547 | 0.94 | 2018 and 2019 |
| 2 | Cameroon | 15,333 | 1.91 | 2017 to 2019 |
| 3 | Egypt | 9,366 | 1.17 | 2017 to 2020 |
| 4 | Eswatini | 3,957 | 0.49 | 2016 to 2018 |
| 5 | Ethiopia | 2,121 | 0.26 | 2016 to 2020 |
| 6 | Gabon | 5,373 | 0.67 | 2016 to 2018 |
| 7 | Ghana | 6,986 | 0.87 | 2016 to 2018 |
| 8 | Kenya | 6,272 | 0.78 | 2016 to 2020 |
| 9 | Madagascar | 16,573 | 2.07 | 2019 |
| 10 | Malawi | 8,264 | 1.03 | 2016 to 2020 |
| 11 | Mali | 2,048 | 0.26 | 2019 |
| 12 | Mozambique | 321 | 0.04 | 2019 |
| 13 | Nigeria | 13,941 | 1.74 | 2016 to 2018 |
| 14 | Senegal | 11,507 | 1.43 | 2016 to 2018 |
| 15 | Sierra Leone | 108 | 0.01 | 2016 to 2020 |
| 16 | South Africa | 488,511 | 60.87 | 2017 to 2020 |
| 17 | Sudan | 979 | 0.12 | 2017 to 2019 |
| 18 | Tanzania | 6,358 | 0.79 | 2016 to 2018 |
| 19 | Tunisia | 158,960 | 19.81 | 2017 to 2020 |
| 20 | Uganda | 25,316 | 3.15 | 2016 to 2020 |
| 21 | Zambia | 10,160 | 1.27 | 2016 to 2020 |
| 22 | Zimbabwe | 2,553 | 0.32 | 2016 to 2018 |
|  | **Total** | **802,554** |  |  |
|  | **Mean** | **36,480** |  |  |
|  | **Standard deviation** | **106,148** |  |  |
|  | **Median** | **6,629** |  |  |
|  | **IQR** | **10,296.75** |  |  |

To provide a more robust description of the highly skewed distribution of isolate counts across countries, additional summary statistics including the median and interquartile range (IQR) were calculated for both the public and private surveillance datasets. For the public datasets, the median number of isolates per country was 6,629 with an interquartile range (IQR) of 10,296.75, whereas the private datasets demonstrated a median of 315 isolates and an IQR of 699.5. These statistics complement the mean and standard deviation by providing measures that are less influenced by extreme values and highly concentrated reporting from a small number of countries, particularly South Africa. The inclusion of the median and IQR therefore provides a clearer representation of the uneven distribution and substantial disparities in AMR surveillance contributions across African countries.

The data in Table 2 shows the number and percentage of isolates from private datasets (ATLAS, DREAM, GEARS, SIDERO-WT, and SOAR) across 13 countries from 2004 to 2022. South Africa again reported the highest number of isolates, with 20,103, accounting for 63.20% of the total, followed by Morocco with 4,409 isolates (13.86%) and Nigeria with 3,720 isolates (11.69%). Other countries, such as Malawi and Uganda, contributed far fewer isolates, at just 59 (0.19%) and 123 (0.39%), respectively. The overall mean number of isolates per country was 2,447, with a standard deviation of 5,490, indicating considerable variation in the data contributed by different countries, thus, similar to results reported by. ^6,7,8^

**Table 2:** The table summarizes AMR data collected from private databases^12^ (ATLAS, DREAM, GEARS, SIDERO-WT, and SOUR) across 13 African countries. It includes the number of isolates for each country and provides the mean and standard deviation of AMR data from 2004 to 2024, as detailed in the table.

| No | Country | Total isolates | Percentage | Year |
| --- | --- | --- | --- | --- |
| 1 | Cameroon | 577 | 1.81 | 2020 to 2022 |
| 2 | Egypt | 315 | 0.99 | 2011 to 2013 |
| 3 | Ghana | 85 | 0.27 | 2017 to 2022 |
| 4 | Ivory Coast | 521 | 1.64 | 2020 to 2022 |
| 5 | Kenya | 736 | 2.31 | 2013 to 2022 |
| 6 | Malawi | 59 | 0.19 | 2021 and 2022 |
| 7 | Mauritius | 119 | 0.37 | 2009 |
| 8 | Morocco | 4,409 | 13.86 | 2004 to 2022 |
| 9 | Namibia | 139 | 0.44 | 2008 and 2009 |
| 10 | Nigeria | 3,720 | 11.69 | 2013 to 2022 |
| 11 | South Africa | 20,103 | 63.2 | 2004 to 2009, 2011 to 2022 |
| 12 | Tunisia | 905 | 2.84 | 2010 to 2017 |
| 13 | Uganda | 123 | 0.39 | 2021 and 2022 |
|  | **Total** | **31,811** |  |  |
|  | **Mean** | **2,447** |  |  |
|  | **Standard deviation** | **5,490** |  |  |
|  | **Median** | **315** |  |  |
|  | **IQR** | **699.5** |  |  |

Table 3 summarizes the inequality and concentration metrics for the distribution of AMR isolates across countries in both the public and private datasets. The public datasets (2016– 2020) demonstrated substantial inequality in country-level isolate contributions, with a Gini coefficient of 0.79 and a Herfindahl–Hirschman Index (HHI) of 0.41, indicating that AMR surveillance data were highly concentrated among a few countries. South Africa alone contributed 60.87% of all public isolates, while the top three contributing countries accounted for 82.42% of the total public dataset. Similarly, the private datasets (2004–2022) also showed marked concentration, with a Gini coefficient of 0.73 and an HHI of 0.46. In the private datasets, South Africa contributed 63.20% of isolates, and the top three contributing countries accounted for 88.75% of all isolates. The Lorenz curves (Supplementary Fig. S1^40^) further illustrated strong deviation from equal distribution in both datasets, confirming substantial imbalance in country-level AMR surveillance contributions across Africa.

**Table 3:** The table displays the results of separate Chi-Square tests of proportions conducted to assess whether the distribution of isolates differed significantly across countries within each dataset. The tests were performed independently for the public database^9,11^ (2016 to 2020) and the private database^12^ (2004 to 2022) to evaluate any significant variations in the distribution of isolates within each dataset.

| Dataset | Total Isolates | Number of Countries | Gini Coefficient | Shannon Entropy | Normalized Entropy | HHI | Top Country Share (%) | Top 3 Country Share (%) |
| --- | --- | --- | --- | --- | --- | --- | --- | --- |
| Public Databases (GLASS and MAAP) | 802,554 | 22 | 0.79 | 1.47 | 0.48 | 0.41 | 60.87 | 82.42 |
| Private Databases (ATLAS, DREAM, GEARS, SIDERO- | 31,811 | 13 | 0.73 | 1.28 | 0.50 | 0.46 | 63.20 | 88.75 |
| WT,<br>SOAR) |  |  |  |  |  |  |  |  |

### Published AMR research data

Table 4 summarises 14 published AMR research studies from 2010 to 2024, showing data contributions from various African countries.^15,16^ Recent contributors include Algeria^19^, South Africa^33^, and Uganda^35^, with publications in 2023 and 2024. Angola has multiple studies from 2015, 2017, and 2018.^20,21,22,23^ Other countries with recent publications are Benin (2022)^24^, Ethiopia (2021)^29^, and Ghana (2020).^30^ The Central African Republic^25^ and Eritrea^28^ each published in 2019, while the Democratic Republic of Congo^26^ and Egypt^25^ contributed in 2018 and 2016, respectively. Mauritius^31^ and Mozambique^32^ have studies from 2015 and 2017, and Tanzania’s^34^ research dates back to 2010 (Supplementary Table S5). This reflects active and diverse AMR research across Africa over the past decade.

**Table 4:** The table presents a summary AMR research data from review papers^15,16^, covering 14 African countries. It details the year of publication and references for studies conducted between 2010 and 2024.

| No | Country | Year of publication | Reference |
| --- | --- | --- | --- |
| 1 | Algeria | 2023 | 19 |
| 2 | Angola | 2015, 2017, and 2018 | 20 – 23 |
| 3 | Benin | 2022 | 24 |
| 4 | Central African Republic | 2019 | 25 |
| 5 | Democratic Republic of Congo | 2018 | 26 |
| 6 | Egypt | 2016 | 27 |
| 7 | Eritrea | 2019 | 28 |
| 8 | Ethiopia | 2021 | 29 |
| 9 | Ghana | 2020 | 30 |
| 10 | Mauritius | 2015 | 31 |
| 11 | Mozambique | 2017 | 32 |
| 12 | South Africa | 2024 | 33 |
| 13 | Tanzania | 2010 | 34 |
| 14 | Uganda | 2024 | 35 |

### Interactive GIS dashboard of private and public databases

The interactive GIS dashboard in Fig 1 presents an extensive overview of 834,365 isolates from both private and public datasets. It distinguishes between private sources like ATLAS, DREAM, GEARS, SIDERO-WT, SOAR and public sources such as MAAP and GLASS. The dashboard illustrates the distribution of these isolates across different countries and years. The bar charts show that private datasets have been collected over a longer period, starting in 2004, while public datasets began collection more recently, in 2016. The map on the right visually displays the data collection locations across Africa, highlighting the countries involved. The timeline at the bottom spans the data collection years, from 2004 to 2022. The dashboard can be accessed at.^41^.

To illustrate how to use the dashboard, consider Cameroon as an example in Fig 2. The private and public dashboard presents a detailed summary of various data points, including country, data source, classification, and year. Users can filter the data based on these parameters. For example, Fig 2A shows that Cameroon has data only from the MAAP public database, with a total of 15,333 isolates collected from 2017 to 2019, but no data from GLASS. In contrast, the private database records only 577 isolates from Cameroon, collected by ATLAS between 2020 and 2022 (Fig 2B).

**Fig 2:** Interactive GIS dashboard overview of antimicrobial resistance data for Cameroon. Figure 2A displays public datasets^9,11^ (MAAP and GLASS) and their distribution across years and geographical locations in Cameroon. The MAAP dataset included 15,333 isolates collected between 2017 and 2019, whereas no isolates were available from the GLASS dataset. Figure 2B presents private dataset^12^ information from ATLAS, showing 577 isolates collected between 2020 and 2022. Both panels include geographical visualization of country-level data collection locations using country centroid coordinates.

### Interactive GIS dashboard of ATLAS data

Fig. 3 presents an interactive dashboard developed using private antimicrobial resistance surveillance data obtained from the ATLAS database, covering isolates collected between 2004 and 2022 across 13 African countries. The dashboard provides a comprehensive overview of AMR patterns and associated metadata, including bacterial species, country of origin, sample source, isolate identifiers, phenotype classifications, year of collection, and antibiotic susceptibility profiles (Fig. 3A). The platform allows users to dynamically filter the dataset by species, country, phenotype, year, and antibiotic, enabling targeted exploration of resistance trends across countries and bacterial pathogens. The integrated GIS component further supports spatial visualization of isolate distribution across Africa, while synchronized charts and tables facilitate interactive analysis of antibiotic response patterns.

**Fig 3:** ATLAS dashboard overview from 2004 to 2022. (A) Comprehensive dashboard view showing datasets from 13 African countries, including bacterial species, sample sources, country distribution, antibiotic resistance profiles, temporal trends, and isolate metadata. (B) Interactive dashboard filtered for *Escherichia coli*, illustrating resistance and susceptibility patterns across multiple antibiotics, together with geographic distribution and country-level isolate counts. (C) Interactive dashboard filtered for *Pseudomonas aeruginosa*, a WHO 2024 bacterial priority pathogen, showing antibiotic resistance profiles, geographic distribution, and synchronized dashboard visualizations derived from ATLAS surveillance data.

As illustrated in Fig. 3B, filtering the dashboard for *E. coli* demonstrates distinct resistance and susceptibility patterns across multiple antibiotics. High levels of resistance were observed for Ampicillin, Levofloxacin, and Ceftriaxone, as indicated by the prominent red resistance bars. In contrast, substantial susceptibility was observed for Amikacin and Imipenem, where the green susceptibility bars dominated the distribution. Intermediate responses were also detected for several antibiotics, including Cefepime and Levofloxacin, reflecting variability in antimicrobial response profiles within *E. coli* isolates across the participating countries. The geographic distribution panel further showed that the majority of *E. coli* isolates originated from South Africa, followed by Nigeria and Morocco, highlighting differences in isolate contribution and surveillance intensity across countries.

Fig. 3C highlights resistance patterns for *Pseudomonas aeruginosa*, one of the pathogens included in the 2024 WHO Bacterial Priority Pathogens List (BPPL). The dashboard revealed elevated resistance levels to Cefepime, Levofloxacin, and Imipenem among *P. aeruginosa* isolates, while susceptibility remained relatively higher for Amikacin. A substantial proportion of isolates also contained “No Information” categories for some antibiotics, reflecting variability in antibiotic testing coverage across surveillance sites and years. Spatial visualization showed that most *P. aeruginosa* isolates were reported from South Africa, Morocco, and Nigeria, consistent with broader isolate distribution patterns observed in the ATLAS dataset.

In summary, the ATLAS dashboard demonstrates how integrated GIS-enabled visualization platforms can support exploration of AMR trends across bacterial species, antibiotics, countries, and years. By combining interactive filtering, synchronized visualizations, and spatial analysis, the dashboard provides a flexible framework for identifying resistance hotspots, examining antibiotic-specific response patterns, and supporting comparative analysis of AMR surveillance data across Africa.

### Antimicrobial resistance heatmap across African countries and antibiotics

The AMR heatmap developed in this study effectively visualises resistance patterns across African countries and antibiotics using aggregated resistance summaries from integrated public and private surveillance datasets (Supplementary Table 2 and Fig 4). The heatmap enables comparison of resistance frequencies across multiple antibiotic-country combinations, highlighting substantial geographical variability in AMR patterns. Several countries demonstrated elevated resistance frequencies to commonly used antibiotics, while selected reserve antibiotics showed comparatively lower resistance levels in some settings. The heatmap further provides a simplified analytical overview of country-level AMR trends and complements the interactive GIS dashboards by enabling rapid identification of resistance hotspots across surveillance systems. Supplementary Fig S1 and Fig S2 further illustrate pathogen-antibiotic and country-pathogen resistance patterns across the integrated datasets.

**Fig 4:** Interactive dashboard developed using R Shiny, visualizing published AMR research papers^15,16^ across 14 African countries listed in Table 3 from 2010 to 2024. Users can select countries and species to explore antibiotic resistance data, view highlighted resistance percentages, and examine sample sources. The dashboard also allows downloading results in CSV format.

### Interactive dashboard of published AMR research data

The interactive dashboard created in this study effectively compiles and visualises samples of published research papers on AMR data collection across African countries (Fig 5). For instance, the dashboard enables users to select a specific country, Mauritius and species, such as *Salmonella*, to explore antibiotic resistance data. In the “Antibiotic Resistance by Country” section, resistance percentages for various antibiotics are displayed, with notable values like 40%, 80%, and 100% resistance highlighted for certain antibiotics. The “Source of Sample” and “Year of Publication” sections offer additional insights, showing that the samples in the selected studies were obtained from sources such as gut, eggs, and intestine. Users can download the results in a comma-separated values (CSV) format. The dashboard can be accessed at.^43^

**Fig 5:** Interactive dashboard developed using R Shiny, visualizing published AMR research papers^15,16^ across 14 African countries listed in Table 3 from 2010 to 2024. Users can select countries and species to explore antibiotic resistance data, view highlighted resistance percentages, and examine sample sources. The dashboard also allows downloading results in CSV format.

## Discussion

### Private and public databases, and published AMR research data

Our findings collectively reveal substantial inequality and concentration in the distribution of AMR surveillance data across African countries within both public and private datasets, as well as within published AMR research studies. Public datasets, including MAAP and GLASS from 2016 to 2020, demonstrated highly concentrated isolate contributions, with South Africa and Tunisia accounting for the majority of reported isolates. Similarly, private datasets including ATLAS, DREAM, GEARS, SIDERO-WT, and SOAR from 2004 to 2022 were heavily dominated by contributions from South Africa, Morocco, and Nigeria. The inequality and concentration analyses, including Gini coefficients, Herfindahl–Hirschman Index (HHI), Shannon entropy, and Lorenz curves, further demonstrated marked imbalance in country-level surveillance contributions across datasets. Median and interquartile range (IQR) statistics additionally highlighted the highly uneven distribution of isolate reporting across countries, particularly the substantial concentration of AMR data among a limited number of surveillance-active countries. These findings are consistent with previous studies reporting substantial disparities in AMR surveillance representation and reporting capacity across Africa^6–8,16,39^

The substantial disparities observed across surveillance datasets likely reflect multiple structural, operational, and health system-related factors extending beyond biological differences in antimicrobial resistance patterns. Variations in laboratory infrastructure, antimicrobial susceptibility testing capacity, surveillance funding, diagnostic resources, workforce availability, reporting systems, healthcare access, and participation in national and regional surveillance programmes may all contribute to unequal AMR reporting representation across African countries^6–8,16^. Countries with stronger laboratory networks, more established surveillance systems, and greater research investment are therefore more likely to contribute larger and more consistent AMR datasets to both public and private surveillance platforms. In contrast, underrepresented countries may reflect regions with limited surveillance infrastructure, fragmented reporting systems, or reduced access to microbiology services rather than lower AMR burden. Similar surveillance inequalities have previously been reported across low- and middle-income countries where resource limitations and uneven investment in microbiology infrastructure constrain routine AMR monitoring capacity^7,8^. Consequently, the observed concentration patterns should be interpreted primarily as indicators of inequalities in surveillance representation, reporting intensity, and data availability across Africa rather than direct comparative measures of AMR prevalence between countries.

These findings suggest that relying solely on public and private datasets does not provide a comprehensive view of AMR research in Africa. Publications, such as those^6,7,8,16^, have highlighted the insufficiencies in AMR data availability across various African nations. These papers emphasise that many countries, particularly those absent from major datasets, face significant data gaps that hinder effective AMR surveillance and response. Thus, the information from published research data, which includes countries not covered in private and public databases, demonstrates that the current datasets alone do not fully capture the scope of AMR research across the continent. This discrepancy underscores the need for more inclusive data collection efforts to address the evident gaps and provide a more accurate picture of AMR dynamics in Africa.

To address these gaps, it is recommended that future AMR research initiatives focus on expanding data collection and reporting to include underrepresented countries and regions. Enhancing collaboration between public health institutions, private sector entities, and research organisations can improve data coverage and accuracy. Additionally, efforts should be made to integrate and harmonise data from various sources, including published research, to provide a more complete and nuanced understanding of AMR in Africa. Encouraging the publication of research from less represented regions and increasing support for AMR surveillance in these areas will also contribute to a more comprehensive view of AMR trends across the continent.

### Interactive GIS dashboards for public, private, and published AMR datasets

The interactive GIS dashboards developed in this study provide an integrated framework for visualising AMR surveillance data from private databases, public surveillance systems, and published research studies across Africa. Rather than functioning as continuously updating real-time surveillance systems, the dashboards were designed as interactive and updateable platforms that support exploration of temporal, geographical, and surveillance-related AMR patterns across multiple datasets and reporting periods. Similar GIS-based approaches have previously been applied in infectious disease surveillance and public health mapping^13^

The dashboards aggregate AMR information across countries and years while preserving the original structure of the contributing datasets. By integrating private datasets (ATLAS, DREAM, GEARS, SIDERO-WT, and SOAR), public surveillance systems (GLASS and MAAP), and published AMR research studies, the dashboards provide a broader representation of AMR surveillance coverage across Africa than would be possible using any single source independently^9,11,12,15,16^. This integration also highlights important differences in temporal coverage, isolate reporting intensity, country participation, and antibiotic testing practices across surveillance systems.

The use of Cameroon as a case study demonstrates the dashboard’s capability to support targeted exploration of AMR surveillance data by country, source, classification, and reporting year. Such filtering functionality allows users to identify surveillance gaps, differences in reporting coverage, and temporal changes in AMR data availability across settings. This functionality may be particularly useful in regions where AMR surveillance data remain fragmented or inconsistently reported^6,7,8^. The ATLAS-specific dashboard further demonstrates the utility of interactive visualisation for isolate-level AMR exploration across multiple African countries. By enabling filtering by bacterial species, sample source, phenotype, antibiotic, and year, the dashboard supports detailed assessment of resistance profiles and susceptibility patterns. The integration of WHO Bacterial Priority Pathogens List (BPPL) data also supports targeted examination of clinically important resistant pathogens relevant to regional AMR priorities. For published AMR research data, the dashboard provides an additional mechanism for visualising research findings that are often absent from formal surveillance systems. Integrating published studies alongside surveillance databases helps broaden representation of countries and datasets that may otherwise remain underrepresented in continental AMR analyses^6,15,16^. The inclusion of metadata such as publication year, sample source, and bacterial species further supports contextual interpretation of AMR trends across countries and time periods.

### Antimicrobial resistance heatmaps across countries, pathogens, and antibiotics

The heatmap highlights substantial variability in resistance frequencies across countries and antibiotics, demonstrating important differences in antimicrobial susceptibility profiles, isolate reporting intensity, and antibiotic testing practices across surveillance settings. Several commonly used antibiotics, including Ampicillin, Ciprofloxacin, Levofloxacin, and Ceftazidime, demonstrated elevated resistance frequencies across multiple countries, suggesting widespread circulation of resistant bacterial strains. In contrast, reserve antibiotics such as Imipenem, Meropenem, and Amikacin generally demonstrated comparatively lower resistance frequencies in selected settings, although variability remained evident across countries and pathogens.

Supplementary Fig S2A further demonstrates the utility of heatmap visualisation for pathogen-antibiotic resistance exploration within the ATLAS dataset. By summarising resistance frequencies across bacterial species and antibiotics, the figure supports comparative assessment of susceptibility profiles among clinically important pathogens including *E. coli*, *K. pneumoniae*, *A. baumannii, P. aeruginosa*, and *S. aureus*. Several pathogens demonstrated elevated resistance frequencies to multiple antibiotic classes, highlighting multidrug resistance patterns across surveillance settings. Gram-negative organisms, particularly *E. coli, K. pneumoniae, A. baumannii,* and *P. aeruginosa*, exhibited higher resistance frequencies to fluoroquinolones and cephalosporins in several countries. Supplementary Fig S2B extends this analysis by visualising country-pathogen resistance distributions, thereby illustrating geographical variation in pathogen-level resistance patterns across African countries included in the ATLAS dataset. Certain pathogens demonstrated consistently elevated resistance frequencies across multiple countries, whereas others showed more localised resistance patterns or lower reporting intensity. Together, these heatmaps complement the interactive ATLAS dashboard by providing condensed analytical summaries of AMR resistance dynamics and supporting identification of resistance hotspots, pathogen-specific resistance trends, and geographical disparities in antimicrobial susceptibility patterns across African surveillance settings. (Sup 2A is Sup 2B)

### Impacts of the GIS interactive dashboard on policy, research, and public health

The interactive GIS dashboards developed in this study may support decision-making, research, and public health activities by improving access to geographically organised AMR surveillance information across multiple African datasets. For policymakers, the dashboards provide a consolidated framework for exploring surveillance coverage, identifying reporting gaps, and supporting evidence-informed planning and resource allocation. The dashboards may also facilitate collaboration between surveillance programmes, researchers, and public health institutions by improving visibility of available AMR data sources^6,7,8^.

From a practical implementation perspective, the dashboards may provide several operational benefits for ministries of health, national AMR coordinating committees, surveillance officers, laboratory networks, researchers, and clinicians involved in antimicrobial resistance monitoring activities. By integrating data from multiple surveillance systems into a unified visualization framework, the dashboards enable users to identify countries, regions, pathogens, and antibiotic-pathogen combinations with limited reporting coverage or inconsistent surveillance participation. Such functionality may assist ministries of health and national AMR focal points in identifying surveillance gaps, prioritising geographical regions requiring expanded laboratory capacity, and supporting strategic planning within National Action Plans (NAPs) for antimicrobial resistance^9,40^. The dashboards may also support regional AMR coordination initiatives by improving visibility of surveillance coverage across multiple countries and facilitating exploratory comparison of temporal reporting trends across surveillance systems^6,7,8^.

At the surveillance and laboratory level, the dashboards may assist surveillance officers and public health institutions in evaluating reporting completeness, monitoring country-level participation over time, and identifying fluctuations in isolate reporting across pathogens and antibiotics. The integration of spatial visualisation tools further allows rapid identification of geographical disparities in AMR data availability and may support prioritisation of laboratory strengthening activities, specimen referral systems, and surveillance expansion efforts in underrepresented regions^13^. In settings where AMR surveillance resources remain limited, the dashboards may also contribute to evidence-informed allocation of laboratory resources, training programmes, diagnostic infrastructure, and antimicrobial stewardship interventions by highlighting areas with reduced surveillance coverage or elevated resistance reporting frequencies^4,5^.

Several additional public health applications may potentially be supported through the use of these dashboards within national and regional AMR surveillance activities. By integrating country-level reporting summaries, pathogen distributions, temporal trends, and spatial visualisation tools, the dashboards may assist ministries of health and surveillance programmes in prioritising geographical regions with limited surveillance coverage or inconsistent reporting participation. Countries demonstrating persistently low isolate reporting, limited laboratory participation, or reduced pathogen-antibiotic representation may require additional microbiology infrastructure, antimicrobial susceptibility testing capacity, workforce training, and specimen referral support^6,7,8^. The dashboards may therefore provide an exploratory framework for identifying regions where strengthened laboratory and surveillance investments could improve AMR reporting representation and surveillance coordination activities.

Although the dashboards were not designed as direct operational policy tools, the integrated visualisation framework may support translation of AMR surveillance observations into exploratory planning and prioritisation activities. For example, countries demonstrating persistently limited surveillance participation, reduced isolate reporting, or restricted pathogen-antibiotic representation may be identified as potential candidates for expanded laboratory strengthening initiatives, workforce development programmes, antimicrobial susceptibility testing support, or increased surveillance investment. Similarly, visualisation of temporal reporting patterns may help surveillance programmes identify interruptions, declines, or inconsistencies in reporting activity across countries, pathogens, or surveillance systems, thereby supporting future planning for improved surveillance continuity and reporting coordination^6,7,8^.

The dashboards may also support exploratory prioritisation of pathogen-antibiotic combinations that remain underrepresented within existing surveillance systems and may therefore require additional investigation through future AMR studies or targeted surveillance initiatives. Spatial visualisation of resistance summaries and isolate reporting distributions may additionally assist ministries of health, surveillance officers, and public health institutions in identifying geographical areas with elevated multidrug resistance reporting frequencies or limited surveillance representation that could warrant further epidemiological assessment or laboratory capacity strengthening. Importantly, these examples are intended to illustrate how integrated visualisation of fragmented AMR datasets may support broader surveillance interpretation and prioritisation activities rather than provide direct operational recommendations or real-time policy decisions. Consequently, the dashboards should be interpreted primarily as exploratory surveillance-support and analytical visualisation resources that may help inform future AMR surveillance strengthening, data harmonisation, and One Health coordination efforts across African settings^4,7,8^.

The geographical and temporal visualisation capabilities may also support exploratory identification of multidrug resistance hotspots and changing resistance reporting patterns across countries and surveillance periods. For example, heatmap summaries and pathogen-antibiotic resistance visualisations may assist users in identifying countries or pathogens demonstrating elevated resistance frequencies to commonly used antibiotics such as fluoroquinolones, cephalosporins, and β-lactam antibiotics. Similarly, the dashboards may support monitoring of temporal fluctuations in isolate reporting and resistance summaries by enabling comparison of reporting intensity across years, pathogens, and surveillance systems. Although these visualisations do not provide real-time outbreak detection or direct operational surveillance alerts, they may assist public health stakeholders in interpreting broader AMR reporting trends and identifying areas warranting further epidemiological investigation or targeted surveillance strengthening efforts^13^.

The inclusion of WHO Bacterial Priority Pathogens List (BPPL) organisms within the ATLAS-specific dashboard further demonstrates the potential utility of integrated visualisation for exploratory assessment of clinically important resistant pathogens relevant to regional AMR priorities. Through interactive filtering of pathogens, antibiotics, countries, and years, users may examine resistance summaries associated with high-priority multidrug-resistant organisms such as *A. baumannii* and *P. aeruginosa*. These exploratory analyses may support interpretation of regional resistance patterns and contribute to future antimicrobial resistance surveillance harmonisation efforts by improving visibility of fragmented datasets originating from public surveillance systems, private surveillance initiatives, and published research studies^4,7,15,16^.

Importantly, the dashboards were intentionally designed as updateable analytical and visualisation tools rather than direct operational surveillance systems or real-time clinical decision-support platforms. Consequently, the dashboards are not intended to replace existing national AMR surveillance programmes, GLASS reporting systems, laboratory information systems, or formal epidemiological investigations. Instead, they provide an integrated framework for exploratory interpretation of AMR surveillance representation, geographical disparities, resistance reporting patterns, and surveillance coverage across multiple datasets while preserving the original structure and ownership of contributing data sources^9,40^.

Although the dashboards developed in this study demonstrate the feasibility of integrating fragmented AMR datasets into interactive analytical platforms, their long-term sustainability would require substantial institutional, technical, and operational support. Sustainable implementation would depend on the establishment of institutional partnerships involving ministries of health, public health laboratories, academic institutions, surveillance networks, and regional AMR coordinating bodies. Long-term functionality would additionally require dedicated hosting infrastructure, sustainable funding mechanisms, routine data updating workflows, governance frameworks for data sharing and stewardship, and integration with existing national and regional AMR surveillance programmes. User training and technical support would also be essential to ensure effective interpretation and application of dashboard outputs by surveillance officers, laboratory personnel, clinicians, researchers, and policymakers. Consequently, the dashboards presented in this study should currently be interpreted as research prototypes and exploratory analytical tools rather than fully operationalised national AMR surveillance systems.

Future large-scale integration of antimicrobial resistance surveillance data across Africa will require strengthened governance frameworks addressing data ownership, interoperability, ethical oversight, controlled-access mechanisms, and equitable sharing of surveillance resources across institutions and countries. Integration of private-sector datasets alongside public surveillance systems introduces additional governance considerations related to data stewardship, permissions for secondary use, transparency of reporting frameworks, and long-term management of restricted-access resources. Sustainable continental AMR surveillance initiatives will therefore require clearly defined data-sharing agreements, institutional governance structures, harmonised interoperability standards, and coordinated ethical oversight mechanisms capable of supporting secure and equitable multi-sectoral data integration within One Health surveillance environments^4,6,7^.

For clinicians, microbiologists, and researchers, the dashboards provide an exploratory analytical environment for visualising pathogen-antibiotic resistance patterns across countries and years. Users can explore susceptibility trends among clinically important pathogens, assess variations in resistance frequencies across surveillance systems, and identify emerging resistance signals that may warrant further investigation. However, the dashboards were not designed as real-time clinical decision-support systems for direct patient management or antibiotic prescribing decisions. Rather, they function as updateable analytical and visualisation tools intended to support interpretation of AMR surveillance data, hypothesis generation, public health situational awareness, and research exploration across fragmented datasets. The dashboards therefore complement existing surveillance infrastructures such as GLASS and national AMR surveillance systems while preserving the original structure, ownership, and reporting context of the contributing datasets^9^.

An additional practical contribution of the dashboards is their ability to improve visibility of fragmented AMR datasets that are often distributed across disconnected public databases, private surveillance programmes, laboratory reports, and published academic studies. Integrating these multiple evidence streams into a single interactive framework enables broader exploration of AMR reporting patterns and supports more transparent interpretation of surveillance representation across African countries. This integrated approach may facilitate future harmonisation efforts, strengthen collaboration between surveillance stakeholders, and support broader One Health AMR monitoring initiatives involving human, animal, food, and environmental health sectors^4,7,8^. Although the dashboards rely on periodically updated datasets rather than continuously streaming surveillance systems, they provide flexible and scalable analytical platforms that can be updated as additional surveillance data become available.

To improve accessibility and reproducibility of the analytical outputs, supplementary dashboard materials including static dashboard screenshots, exported visualisations, figure panels, and repository documentation were deposited within the Figshare repository associated with this study. These supplementary materials provide alternative mechanisms for accessing and reviewing the dashboard outputs in settings where direct interaction with web-based dashboards may be limited by internet connectivity, computational infrastructure, or digital access constraints. Future work may further improve accessibility and usability through the development of video screencasts, interactive tutorials, end-user documentation, and structured training materials designed to support broader adoption and interpretation of dashboard outputs across diverse public health and surveillance environments.

The dashboards may also be influenced by visibility bias, whereby countries contributing larger and more consistent datasets naturally appear more prominent within the visualisations and analytical summaries. Importantly, greater visibility within the dashboards should not be interpreted as indicating that AMR is less important or less severe in underrepresented countries. Rather, lower representation frequently reflects limited surveillance coverage, reduced laboratory capacity, inconsistent reporting systems, and weaker integration into regional and global AMR surveillance networks. Consequently, countries with fewer reported isolates may in fact represent regions with substantial unmet surveillance needs rather than lower AMR burden. The dashboards should therefore be interpreted as tools that reveal inequalities in AMR surveillance representation and reporting capacity across Africa, while also highlighting regions where strengthened surveillance investment and improved data integration may be most urgently required. By explicitly visualising these disparities, the dashboards may help support more equitable prioritisation of future surveillance strengthening initiatives and broader One Health AMR capacity-building efforts across the continent.4,7,8

For researchers, the dashboards support data exploration across multiple surveillance systems and published studies, enabling identification of underrepresented regions, organisms, and antibiotic-pathogen combinations requiring further investigation. The integration of multiple AMR evidence streams within a single framework may also support future harmonisation and comparative surveillance studies^15,16,17^. For public health stakeholders, the dashboards provide interactive visualisation tools for examining temporal and spatial patterns in AMR reporting across Africa. Although the dashboards are based on periodically compiled datasets rather than continuously streaming surveillance systems, they provide updateable analytical frameworks that can support ongoing surveillance reporting and AMR data interpretation. In addition, the dashboards may contribute to AMR awareness and communication by presenting complex surveillance data in a more accessible and interpretable format^13^.

### Limitations of the study

Several limitations should be considered when interpreting the findings and dashboards presented in this study. First, the integrated AMR datasets originated from multiple surveillance systems and published studies that differed substantially in surveillance duration, reporting intensity, laboratory participation, isolate selection procedures, diagnostic capacity, and antibiotic testing practices. Consequently, direct comparisons between public and private datasets should be interpreted cautiously, as differences in isolate counts may reflect variations in surveillance coverage and reporting infrastructure rather than true differences in AMR burden across countries. Second, the dashboards were developed using datasets extracted at specific time points and therefore do not represent continuously updating real-time surveillance systems. Instead, the dashboards function as interactive and updateable analytical platforms that can support exploration of AMR surveillance data across countries and years. Although this approach enables integrated visualisation of multiple datasets, temporal gaps and reporting inconsistencies remain important limitations.

Third limitation relates to the substantial heterogeneity and potential mismatches between the contributing datasets. Public surveillance systems, private-sector datasets, and published AMR studies differed considerably in surveillance scope, laboratory methodologies, antimicrobial susceptibility testing standards, isolate selection procedures, reporting intensity, metadata completeness, and temporal coverage. These differences may introduce reporting bias, inconsistencies in resistance interpretation, and unequal representation of pathogens, antibiotics, and countries across datasets. Furthermore, temporal mismatches between surveillance periods and variability in data availability across countries may influence the comparability of resistance summaries and geographical visualisations presented within the dashboards. Countries with stronger surveillance systems and more sustained reporting participation may therefore appear more prominent within the dashboards, potentially contributing to visibility bias in the interpretation of surveillance representation across Africa. Consequently, the dashboards should be interpreted primarily as exploratory surveillance-support and visualisation tools designed to improve visibility of fragmented AMR datasets rather than definitive comparative epidemiological surveillance systems.

Fourth, several African countries remain underrepresented or absent from both public and private AMR surveillance databases, highlighting persistent continental gaps in AMR reporting and laboratory infrastructure. Published research studies partially addressed some of these gaps; however, published datasets also varied in study design, reporting standards, bacterial species investigated, and antimicrobial susceptibility testing methodologies. In addition, the study did not standardise or normalise isolate counts across countries because consistent metadata regarding reporting laboratories, population denominators, and surveillance intensity were not uniformly available across all datasets. As a result, isolate counts presented in the dashboards should not be interpreted as direct estimates of AMR prevalence or disease burden. Rather, they primarily reflect surveillance representation and reporting availability across datasets and countries. While dashboards provide efficient visualisation and exploratory analysis of complex AMR data, their utility depends on digital access, internet connectivity, and user familiarity with interactive analytical tools. These factors may limit accessibility in some low-resource settings. Nevertheless, the combination of interactive dashboards and supporting tabular summaries provides complementary approaches for communicating AMR surveillance findings across diverse audiences.

Accessibility of interactive dashboard platforms may remain challenging in some low-resource healthcare and surveillance environments where reliable internet connectivity, computational infrastructure, and digital access remain inconsistent. Although the dashboards were designed to support flexible exploration of AMR surveillance data, variability in digital infrastructure and user familiarity with interactive analytical platforms may influence usability and accessibility across different settings. Potential future approaches to improve inclusivity may include development of simplified dashboard interfaces, downloadable static reports, offline-compatible analytical outputs, mobile-friendly visualisation formats, and integration with existing local surveillance reporting systems. Such approaches may help improve accessibility of AMR surveillance information across diverse operational and resource settings within Africa.

An additional limitation of this study is that formal usability testing and stakeholder validation were not conducted during the initial development of the dashboards. Although the dashboards were designed to support exploratory visualisation and interpretation of AMR surveillance data across Africa, their practicality, accessibility, and operational usefulness for ministries of health, surveillance officers, clinicians, policymakers, laboratory personnel, and other public health stakeholders were not systematically evaluated within this study. Consequently, the dashboards should currently be interpreted as pilot research and visualisation frameworks intended primarily to demonstrate the feasibility of integrating fragmented AMR datasets from multiple public, private, and published surveillance sources across African countries.

Another important limitation is that long-term sustainability, maintenance, hosting responsibilities, and operational governance frameworks for the dashboards were not formally established within the scope of this initial study. The dashboards were developed primarily as pilot research and visualisation frameworks to demonstrate the feasibility of integrating fragmented AMR datasets across Africa rather than as fully operational national surveillance platforms. Consequently, sustainable long-term implementation would require clearly defined institutional ownership, dedicated hosting infrastructure, sustainable financial support, routine mechanisms for data updating and quality assurance, user training programmes, and governance structures for data access, stewardship, and interoperability with existing AMR surveillance systems. Without such long-term institutional and operational support, maintaining regularly updated and accessible dashboard platforms across diverse surveillance environments may remain challenging.

Potential risk of misinterpretation or oversimplification of dashboard findings by non-expert users is another limitation. Because the dashboards integrate heterogeneous datasets originating from multiple surveillance systems with differing laboratory methodologies, surveillance intensities, reporting standards, and isolate selection procedures, visual patterns observed within the dashboards should not be interpreted as definitive measures of national AMR burden, healthcare performance, or direct comparative country rankings. Simplified interpretation of resistance heatmaps, isolate counts, or country-level reporting frequencies without appropriate epidemiological and methodological context could potentially result in inappropriate conclusions, oversimplified policy interpretations, or disproportionate prioritisation decisions. Ethical considerations related to data ownership, restricted-access surveillance resources, and cross-institutional data sharing also remain important considerations for future large-scale AMR integration initiatives. Appropriate user training, governance frameworks, epidemiological guidance, and stakeholder engagement will therefore be essential to support responsible interpretation, communication, and application of dashboard outputs within public health, surveillance, research, and policy environments.

Formal dashboard validation and structured usability assessment involving ministries of health, surveillance officers, clinicians, laboratory personnel, policymakers, and other end-users were not performed during this initial study. Consequently, the dashboards should currently be interpreted as exploratory research and visualisation prototypes intended primarily to demonstrate the feasibility of integrating fragmented AMR datasets across Africa. Future work should therefore include stakeholder co-design approaches, usability studies, pilot implementation assessments, clinician evaluation, public health stakeholder feedback, and policy-oriented validation exercises to improve interpretability, accessibility, operational relevance, and long-term adoption within routine AMR surveillance workflows.

Future large-scale integration of antimicrobial resistance surveillance data across Africa will require strengthened governance frameworks addressing data ownership, interoperability, ethical oversight, controlled-access mechanisms, and equitable sharing of surveillance resources across institutions and countries. Integration of private-sector datasets alongside public surveillance systems introduces additional governance considerations related to data stewardship, permissions for secondary use, transparency of reporting frameworks, and long-term management of restricted-access resources. Sustainable continental AMR surveillance initiatives will therefore require clearly defined data-sharing agreements, institutional governance structures, harmonised interoperability standards, and coordinated ethical oversight mechanisms capable of supporting secure and equitable multi-sectoral data integration within One Health surveillance environments^4,6,7^.

Future work should therefore include stakeholder co-design approaches, usability assessments, implementation evaluation studies, structured user feedback exercises, and broader policy engagement activities to improve dashboard functionality, interpretability, and long-term adoption within routine AMR surveillance workflows. In addition, country-level validation exercises involving national AMR programmes, public health laboratories, surveillance officers, and regional stakeholders would help assess the operational relevance, accessibility, and contextual applicability of the dashboards across diverse surveillance environments. Such collaborative evaluation processes may further support refinement of visualisation features, prioritisation of user needs, improvement of reporting workflows, and development of more sustainable and interoperable AMR surveillance platforms aligned with national and regional public health priorities^4,6,8^.

## Conclusion

The integration of AMR data from private surveillance systems, public databases, and published research studies revealed substantial disparities in AMR reporting coverage across African countries, highlighting important surveillance gaps and unequal data representation across the continent. The interactive GIS dashboards developed in this study provide an integrated and updateable framework for visualising and exploring AMR surveillance data across multiple countries, years, and data sources. By combining private, public, and published AMR datasets within a unified analytical framework, the dashboards improve visibility of surveillance coverage, facilitate identification of reporting gaps, and support interpretation of geographical and temporal AMR patterns across Africa. These tools may support policymakers, researchers, and public health institutions in strengthening surveillance activities, prioritising research needs, and improving evidence-informed decision-making. However, the dashboards rely on periodically compiled datasets and therefore should not be interpreted as continuously updating real-time surveillance systems. In addition, differences in surveillance duration, laboratory participation, and reporting intensity across datasets limit direct comparability between surveillance systems. Future efforts focused on harmonised reporting standards, expanded surveillance participation, and improved data integration across African countries will further strengthen the utility of interactive AMR surveillance platforms.

Beyond improving visibility of fragmented AMR surveillance information, the integrated dashboard framework developed in this study may contribute to broader efforts aimed at strengthening antimicrobial resistance monitoring and surveillance coordination across Africa. By encouraging integration of public, private, and academic AMR datasets, the dashboards support greater awareness of surveillance inequalities, facilitate exploratory identification of reporting gaps, and promote more geographically informed interpretation of resistance patterns across countries and surveillance systems. The dashboards may additionally support future harmonisation initiatives, One Health surveillance approaches, and data-driven AMR monitoring efforts aligned with regional and global public health priorities. Although the dashboards are not intended to function as definitive epidemiological surveillance systems or direct operational policy platforms, they provide exploratory surveillance-support tools designed to improve interpretation, accessibility, and visualisation of fragmented AMR information across Africa. In the longer term, strengthened integration and interpretation of AMR surveillance data may contribute to improved public health preparedness, antimicrobial stewardship planning, and progress toward global antimicrobial resistance control efforts, including the WHO Global Action Plan on Antimicrobial Resistance and Sustainable Development Goal 3 focused on good health and well-being^4^

The findings of this study additionally highlight several important priorities for future antimicrobial resistance surveillance strengthening efforts across Africa. Improved harmonization of AMR surveillance methodologies, expansion of laboratory infrastructure and diagnostic capacity in underrepresented regions, and stronger integration of public, private, and academic AMR datasets will be essential for improving surveillance coverage and comparability across countries. Strengthened data-sharing frameworks, interoperable surveillance systems, and collaborative One Health surveillance approaches may further support coordinated AMR monitoring activities across human, animal, food, and environmental health sectors. In addition, improving accessibility and usability of AMR visualisation tools through stakeholder engagement, user training, implementation studies, and integration with existing surveillance programmes may enhance long-term adoption and operational relevance. Although the dashboards developed in this study are intended primarily as exploratory surveillance-support and analytical visualisation tools rather than definitive epidemiological surveillance systems, they provide a scalable framework for improving interpretation, accessibility, and integration of fragmented AMR surveillance information across Africa while supporting future collaborative surveillance strengthening initiatives aligned with regional and global AMR priorities.

## Acknowledgement

This publication or presentation is based on research using data from Pfizer, GSK, Johnson and Johnson, Venatorx, Shionogi, and Paratek, obtained through https://amr.vivli.org.

## Supporting information

**Figs. 2 and 3**: were generated using analyses derived from the restricted-access Pfizer ATLAS isolate-level dataset. Because the isolate-level ATLAS dataset is subject to controlled-access restrictions, it cannot be publicly redistributed by the authors. However, the deposited derived datasets, including country_year_pathogen_counts.xlsx and resistance_summaries_by_country_year_pathogen.xlsx (**Supplementary Table S2**)**^Error! Reference source not found.^**, the aggregated outputs.

## Data Availability Statement

### Aggregated Datasets

The aggregated datasets are deposited at Figshare repository (https://figshare.com/s/ec72a59e2198485f88b3) with the DOI 10.6084/m9.figshare.32232918. The deposited datasets include:

#### 1) resistance_summaries_by_country_year_pathogen.xlsx (Supplementary Table S1)

**Supplementary Table S1.** contains country-level spatial and temporal summaries of antimicrobial resistance (AMR) surveillance data across Africa. Each record represents the number of bacterial isolates collected within a specific country and year linked to geographic coordinates (x and y) for spatial mapping and GIS visualization. The Country variable identifies the reporting country, while Year captures the surveillance period associated with the isolate counts. The isolate column represents the total number of bacterial isolates reported for each country-year combination. The Classification variable distinguishes surveillance or data source categories, including public and private. The Source variable identifies the originating surveillance platform or database, including ATLAS (Private) and MAAP (Public).

#### 2) country_year_pathogen_counts.xlsx (Supplementary Table 2)

**Supplementary Table 2** contains aggregated counts of bacterial isolates summarized by country, year, and pathogen species derived from the ATLAS (Antimicrobial Testing Leadership and Surveillance) AMR dataset. Each record represents the total number of isolates identified for a specific bacterial species within a given country and year. These aggregated outputs can be used to support temporal analyses, pathogen distribution summaries, and visualization. The dataset provides a reproducible intermediate data layer linking the isolate-level analytical workflow to the reported study outputs.

#### 3) Africa_AMR_Country_Year_Spatial_Data.xlsx (Supplementary Table 3)

**Supplementary Table 3** contains country-level spatial and temporal summaries of antimicrobial resistance (AMR) surveillance data across Africa. Each record represents the number of bacterial isolates collected within a specific country and year, linked to geographic coordinates (x and y) for spatial mapping and GIS visualization. The Country variable identifies the reporting country, while Year captures the surveillance period associated with the isolate counts. The isolate column represents the total number of bacterial isolates reported for each country-year combination. The Classification variable distinguishes the type of surveillance or data source category, such as public or private sector data contributions, enabling comparative analyses between surveillance systems. The Source variable identifies the originating surveillance platform or database, including sources such as ATLAS and MAAP.

### Restricted Dataset

Pfizer ATLAS data through Vivli is restricted dataset. Qualified researchers can obtain the dataset using the following process:

Step-by-Step Process to Obtain Data:

1. Register/Login: Visit searchamr.vivli.org to create an account using a valid email address and complete email verification.
2. Search Datasets: Use the keyword search or browse functions on the AMR Register to identify relevant datasets by country, organism, or antimicrobial agent.
3. Submit Request: Click “REQUEST DATASETS” and provide a brief summary of the proposed research objectives and intended public health relevance.
4. Review Process: Some datasets may require review of the research proposal by Vivli and the original data contributor.
5. Data Access: Once approved, datasets can be downloaded through the “Active” tab on the platform.

Key Information for Access:

- Datasets generally include anonymized isolate-level antimicrobial susceptibility data, including MIC (Minimum Inhibitory Concentration) values and associated metadata.
- Most datasets are available directly through Vivli, except for selected datasets managed through separate contributor portals.
- Users must agree to the terms of use and acknowledge the original data contributors and Vivli in resulting publications.

Accordingly, while the raw isolate-level Pfizer ATLAS data remain under controlled access, the derived aggregated datasets necessary to reproduce the figures, tables, resistance summaries, spatial analyses, and major are publicly available through Figshare.

---

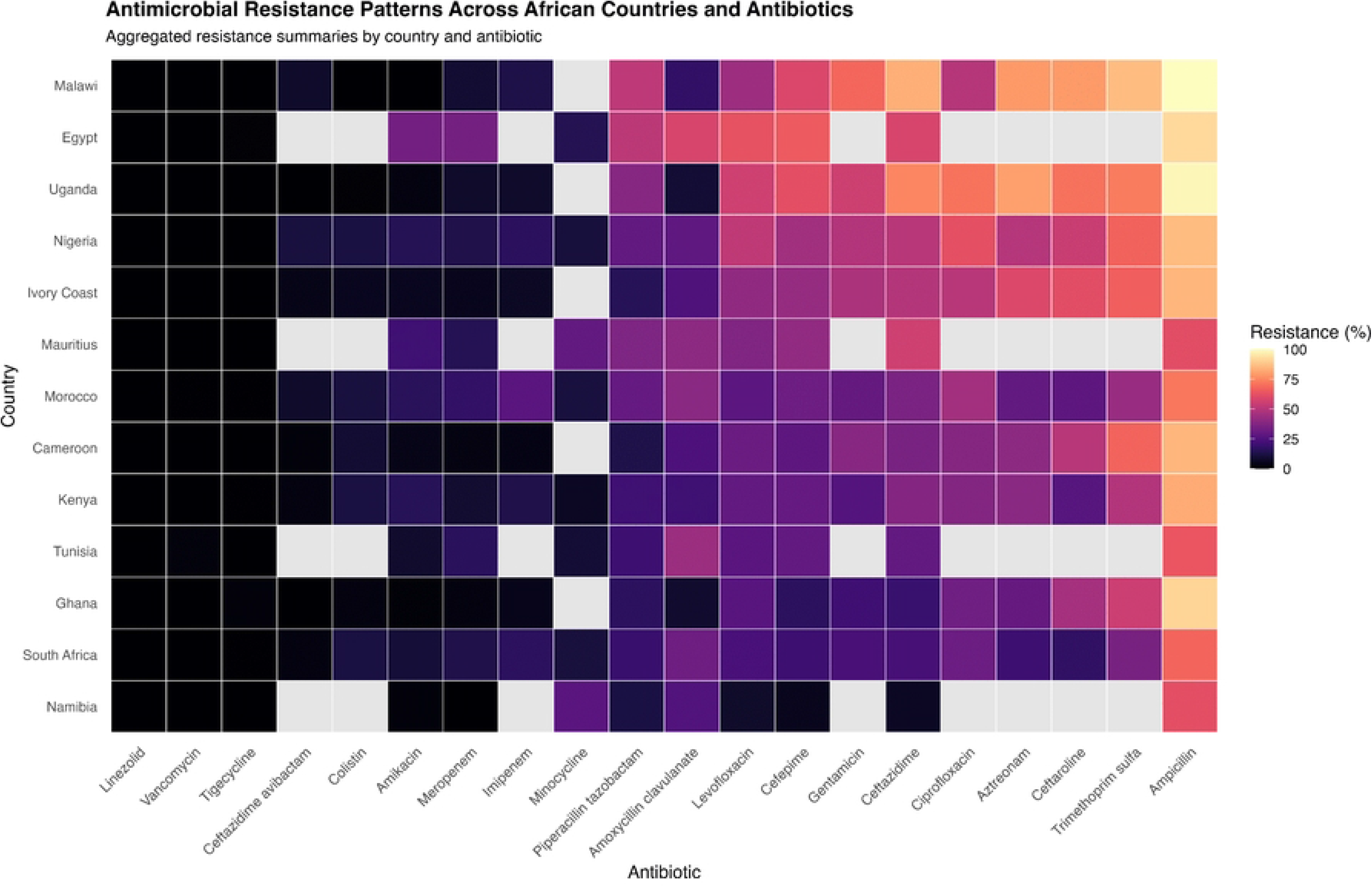

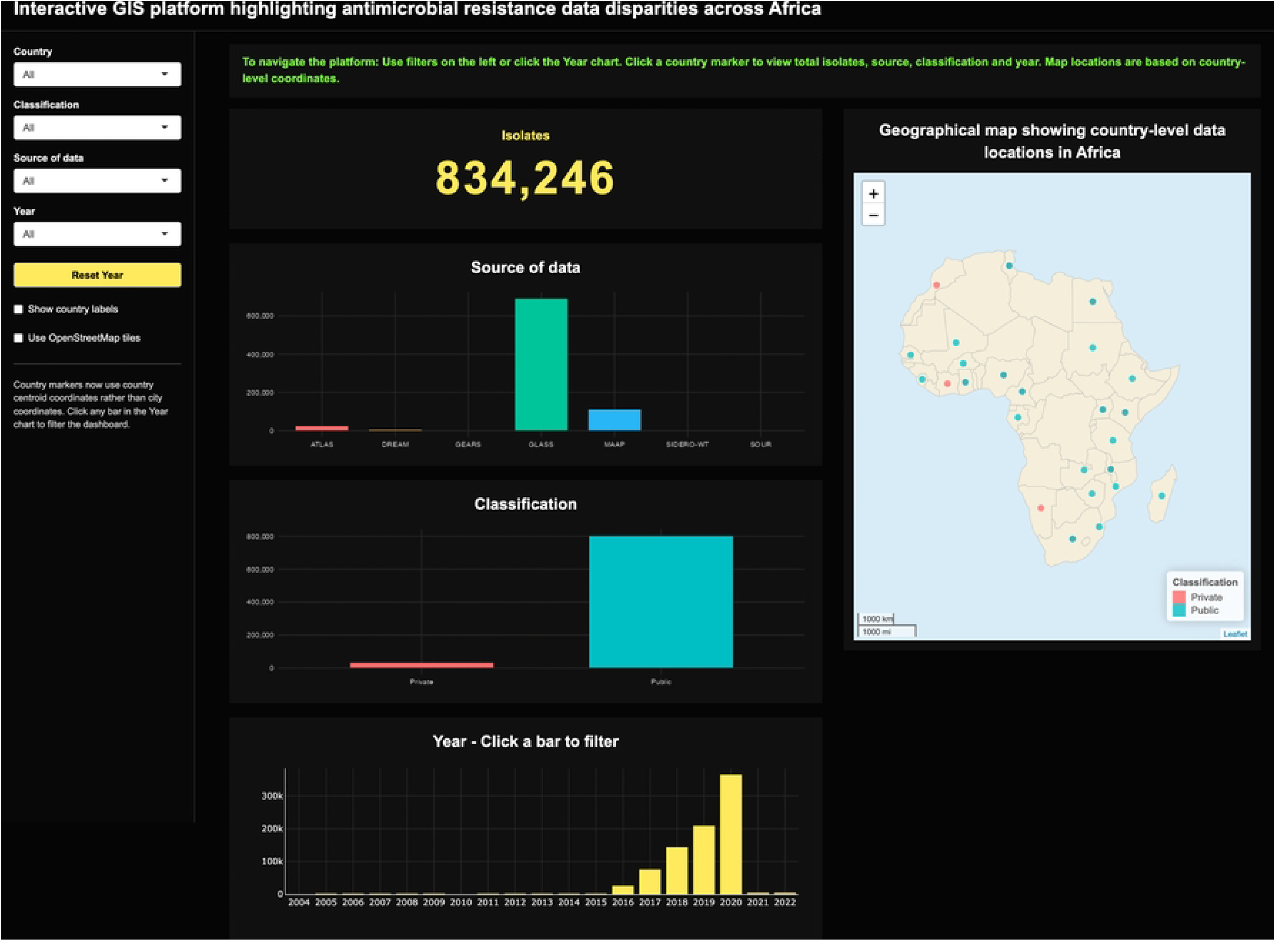

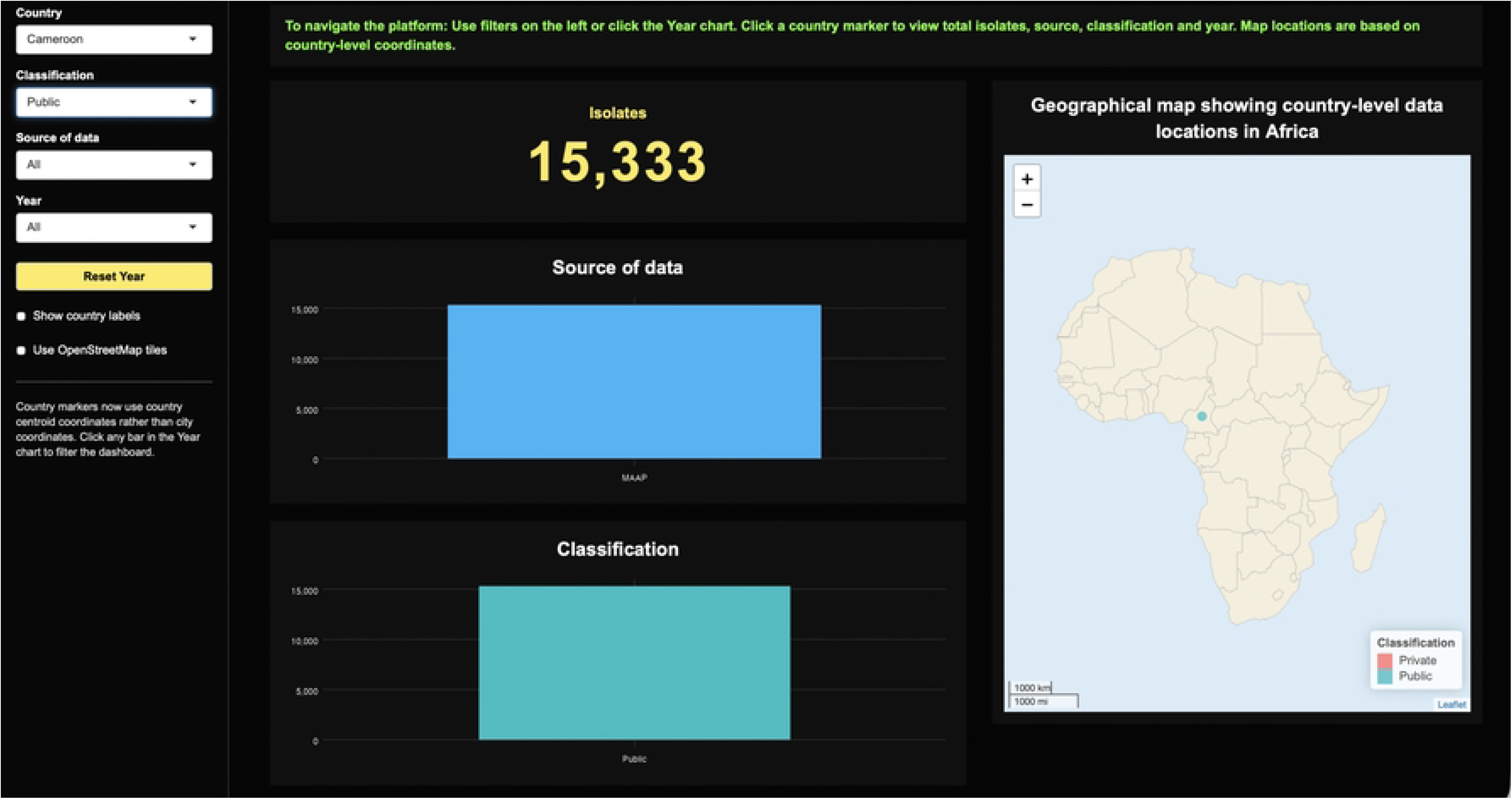

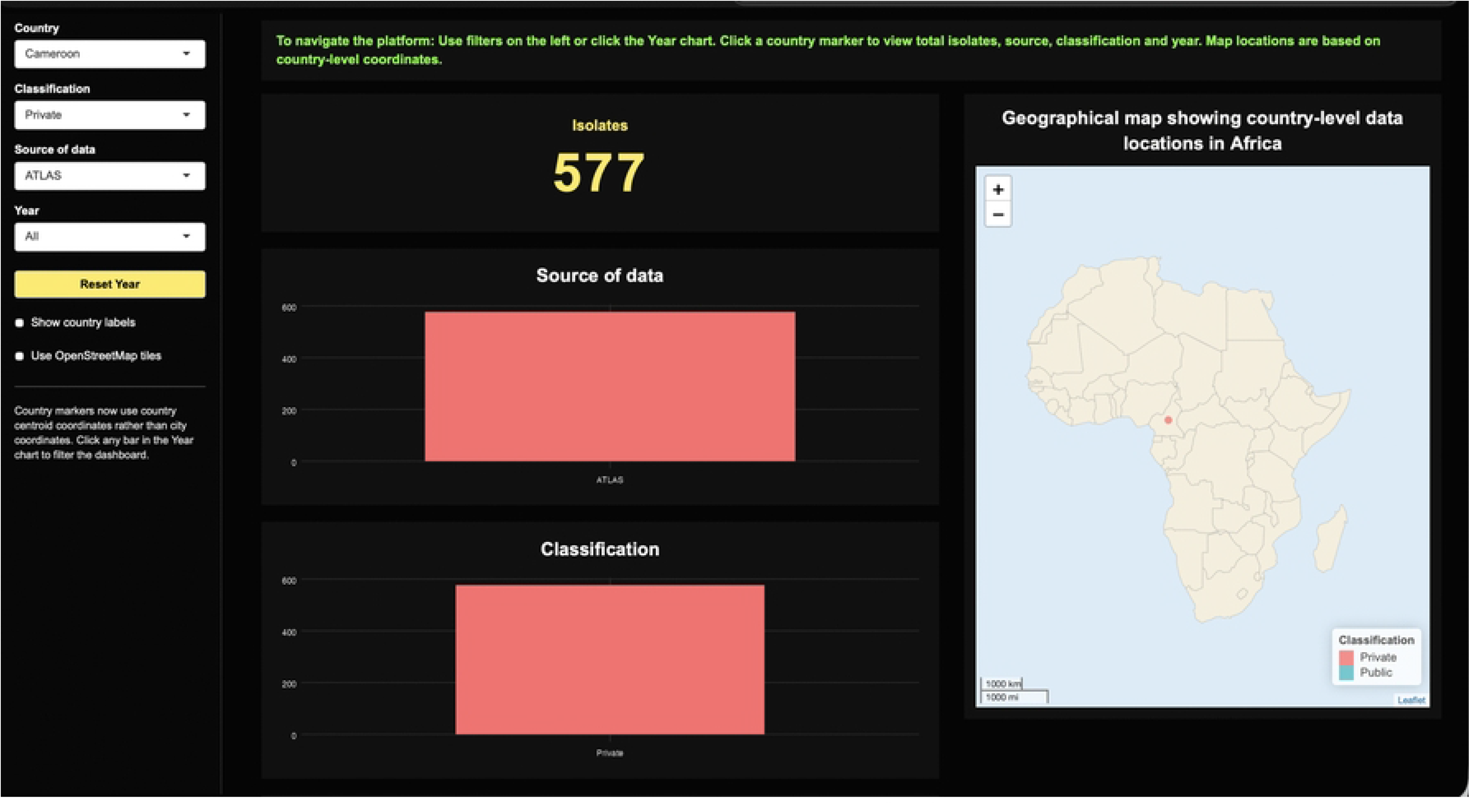

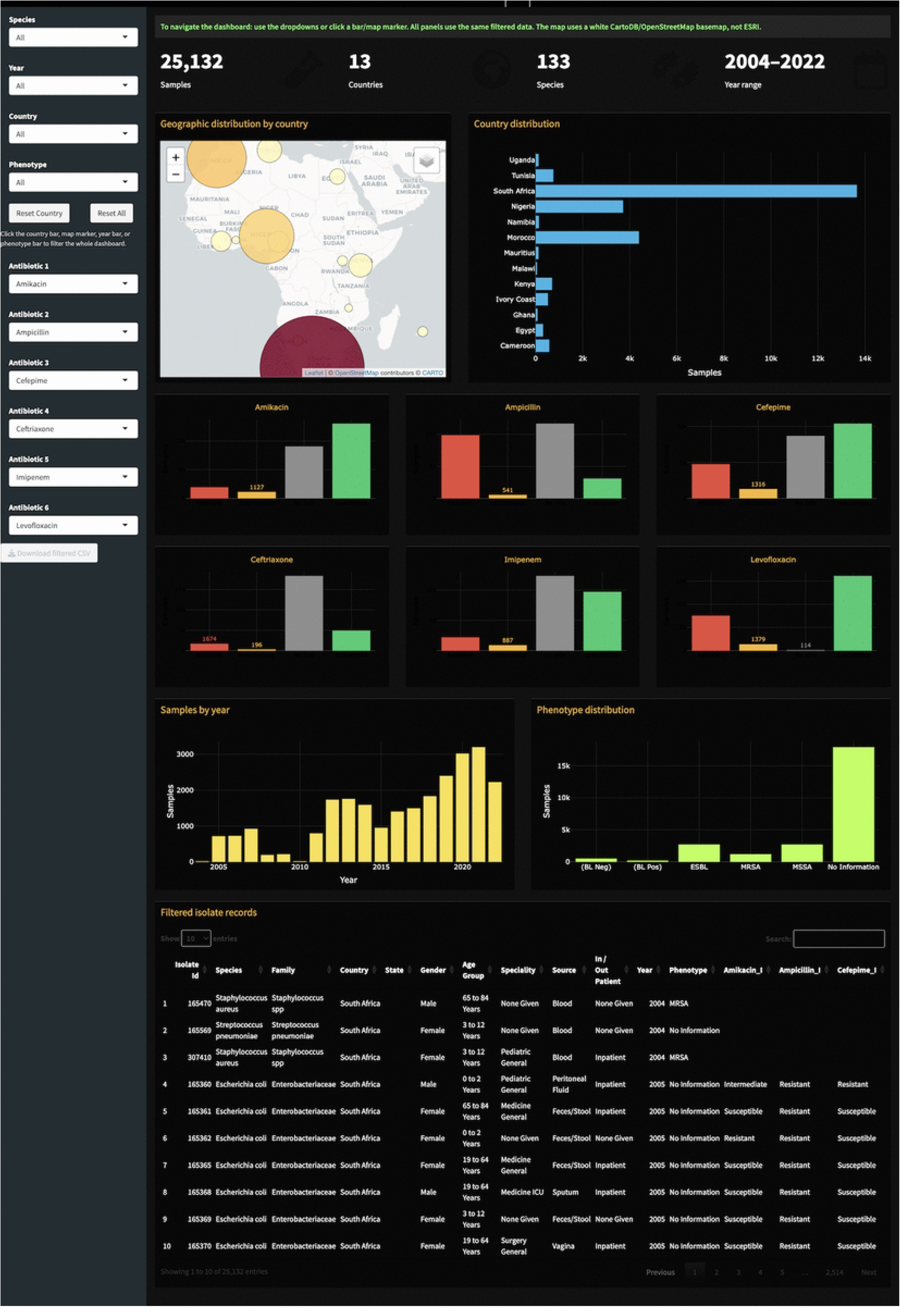

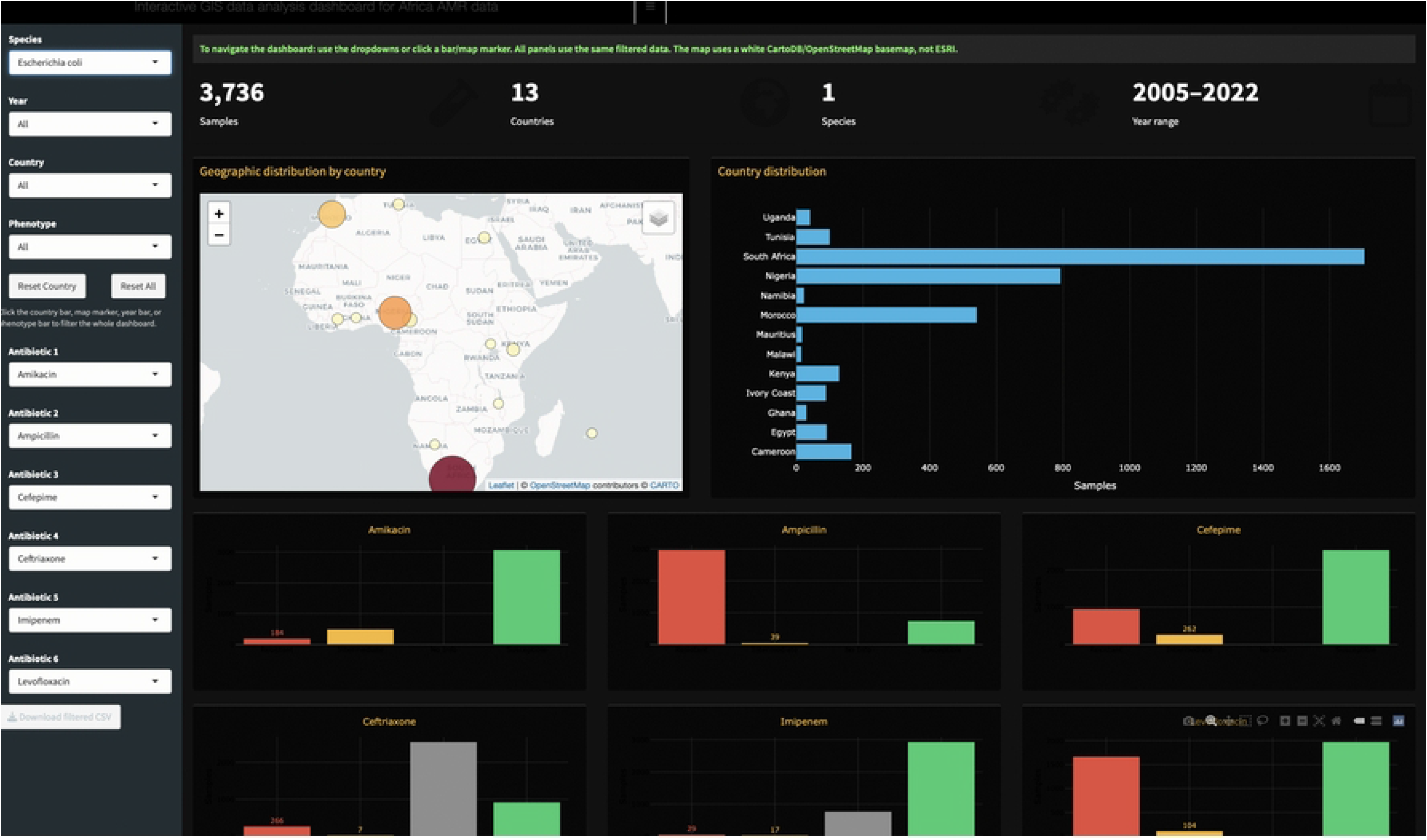

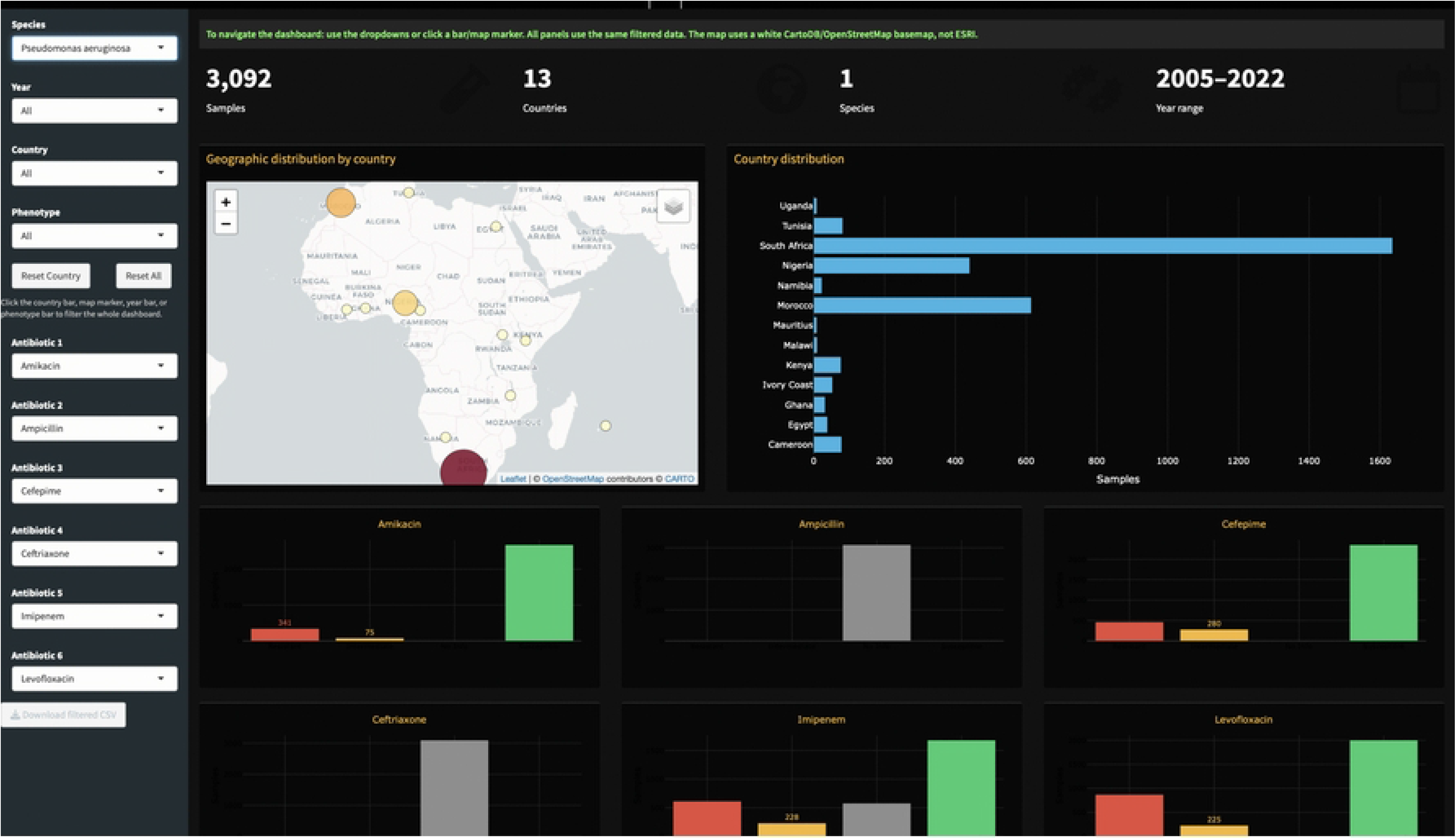

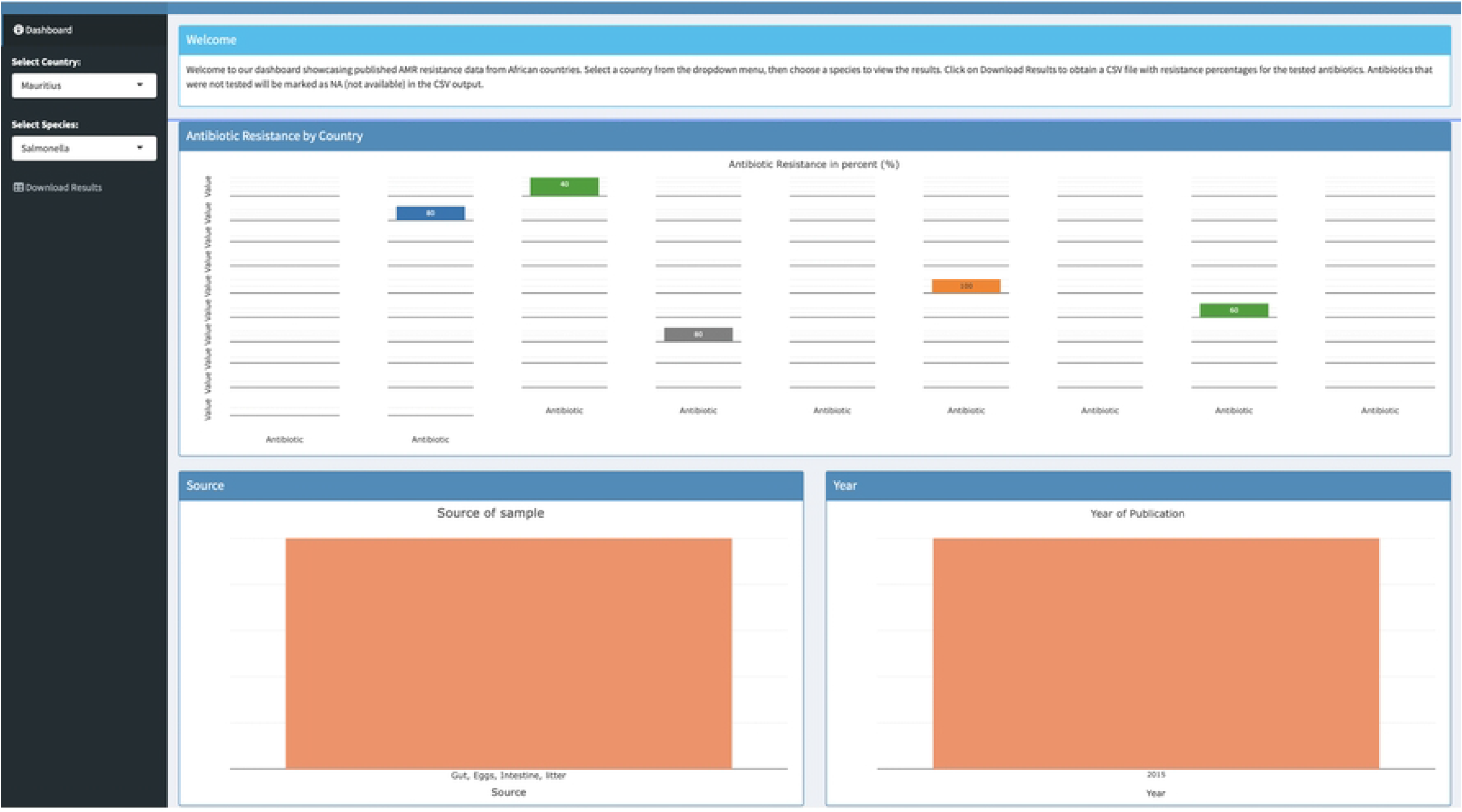

